# Disrupting the Feed-Forward Cycle of RyR1 Ca^2+^ Leak and Oxidative Stress Mitigates Doxorubicin-Induced Skeletal Myopathy

**DOI:** 10.1101/2025.11.12.688073

**Authors:** Lourdes C. Figueroa, Eshwar Tammineni, Pablo Marco-Moreno, Ainara Vallejo-Illaramendi, Adolfo López de Munain, Maialen Sagartzazu, Michael Fill, Carlo Manno

## Abstract

Doxorubicin (DOX), a highly effective and widely used chemotherapeutic agent used to treat various types of cancer. Unfortunately, DOX also has some undesirable and off-target effects, particularly debilitating muscle weakness and fatigue. The mechanism behind this DOX-induced skeletal myotoxicity (DISM) remains unclear. Here, we show that acute DOX exposure, at clinically relevant concentrations, impairs isometric force production and accelerates fatigue in ex vivo murine flexor digitorum brevis (FDB) muscles. Mechanistically, we found that DOX increases the open probability of single RyR1 and disrupts calcium (Ca^2+^)-dependent inactivation (CDI). This results in a persistent sarcoplasmic reticulum (SR) Ca^2+^ leak, elevated basal cytosolic Ca^2+^, and abnormal Ca^2+^ release during action potentials. This abnormal intracellular Ca^2+^ handling ultimately leads to increased mitochondrial reactive oxygen species (ROS) production, which, in turn, exacerbates the functional instability of RyR1. Interestingly, the cytosolic basal Ca^2+^ elevation precedes ROS generation, suggesting that it initiates a destructive cross-talk between Ca^2+^ dysregulation and oxidative stress. Notably, pharmacological stabilization of the RyR1-FKBP12 complex with novel triazole compounds, MP-001 and MP-034, normalizes RyR1 function, Ca^2+^ and ROS homeostasis, as well as muscle force and fatigue resistance. Our findings indicate that DISM is initiated by DOX destabilization of the RyR1-FKBP12 complex (abnormal SR Ca^2+^ leak) and then exacerbated by the Ca-ROS vicious cycle. Limiting RyR1-mediated Ca^2+^ leak with MP-001 represents a promising therapeutic strategy for anti-DISM, aiming to normalize muscle function in patients undergoing DOX chemotherapy.

## Introduction

During excitation-contraction (EC) coupling, a process in which an action potential triggers muscle contraction, the control of calcium (Ca^2+^) release and reuptake is critical for proper muscle function. The functional and structural unit responsible for intracellular Ca^2+^ release in EC coupling was defined as a “couplon” (Stern, 1992). The couplon is mainly composed of dihydropyridine receptors or Ca_V_1.1, a voltage-gated Ca^2+^ channel located on the transverse tubules, and ryanodine receptors (RyR1), which form Ca^2+^ release channels on the sarcoplasmic reticulum (SR). Additionally, other proteins are also involved in this process, including calsequestrin, triadin, junctophilin, and JP-45, among others (Perni *et al*., 2017). Additionally, the presence of endogenous ligand proteins of the RyR1, such as the FK505-binding protein (FKBP12), are important to maintain a stable full-conductance state and therefore normal channel activity (Brillantes *et al*., 1994). Dysfunction within any component of the couplon, whether due to genetic defects or other factors, is defined by our group as “couplonopathy.” Interestingly, the disruption of individual couplon proteins can yield similar phenotypic outcomes, which further highlights the importance of considering these proteins as part of a functional signaling unit (Ríos *et al*., 2015).

Doxorubicin (DOX) is an anthracycline chemotherapeutic agent widely used in the treatment of various types of cancer (Campelj *et al*., 2021). Despite its effectiveness, DOX is associated with a range of adverse effects caused by its off-target interaction in healthy cells, including cardiotoxicity, chemotherapy-induced fatigue, weakness, and skeletal muscle dysfunction, significantly impacting the quality of life of patients undergoing cancer treatment (Morrow *et al*., 2002; Chen *et al*., 2007; Min *et al*., 2015; Siegel *et al*., 2024). The exact nature of Doxorubicin-Induced Skeletal Myotoxicity (DISM) remains unclear (Hydock *et al*., 2011), and it has often been correlated with impaired energy production, oxidative stress, and the activation of cellular proteases, which play a significant role in compromising the ability of muscles to generate force and resist fatigue. It has also been hypothesized that DISM may involve a pathway where DOX directly stimulates the production of reactive oxygen species (ROS), through redox cycling or indirectly via TNF signaling, ultimately promoting mitochondrial dysfunction (Gilliam & Clair, 2011; Gilliam *et al*., 2013*a*; Fabris & MacLean, 2015).

Building upon our understanding of nitro-oxidative stress and emphasizing the role of intramitochondrial Ca^2+^ in ROS production as a contributing factor to DISM (Posterino & Lamb, 2003), observed in various myopathies (Durham *et al*., 2008; Canato *et al*., 2019; Michelucci *et al*., 2021; Reggiani & Marcucci, 2022; Figueroa *et al*., 2023), we hypothesize that DOX’s impact extends further to the production of ROS, suggesting a destructive cross-talk between abnormal Ca^2+^ handling and mitochondrial dysfunction. Specifically, we propose that DOX directly affects Ca^2+^ release units, disrupting Ca^2+^ signaling by systematically increasing the leak of Ca^2+^ from the SR, as previously demonstrated in SR vesicles (Zorzato *et al*., 1985*a*) and in cardiac RyR2 single channel studies (Hanna *et al*., 2014), while simultaneously impairing Ca^2+^ reuptake mechanisms mediated by the SR Ca-ATPase (Norren *et al*., 2009).

Using this previous rationale, we further suggest that the way DOX alters RyR1 activity may be by hindering the channel’s Ca-dependent inactivation (CDI) (Jong *et al*., 1995; Caputo *et al*., 2004; Manno *et al*., 2013*b*), making it more prone to leak Ca^2+^. This, in turn, can ultimately stimulate the production of ROS/RNS, establishing a feed-forward cycle in which elevated oxidative stress further amplifies Ca^2+^ dysregulation. To reinforce this hypothesis, it has been previously found that nitro-oxidative stress disrupts FKBP12-RyR1 interaction, rendering RyR1 channels leaky (Durham *et al*., 2008; Bellinger *et al*., 2009; Aizpurua *et al*., 2021). Consequently, we propose that directly targeting Ca^2+^ dysregulation may represent an effective therapeutic strategy to treat this drug-induced couplonopathy. A similar approach has been applied to mitigate the detrimental effects of DOX on lymphatic node function by blocking the activity of RyR channels (Van *et al*., 2021).

In this study, we employed an ex vivo chemotherapy model, acutely exposing DOX to preparations ranging from single RyR1 channels to whole flexor digitorum brevis (FDB) muscles. We assess the impact of this drug on single-channel properties, cellular Ca^2+^ dynamics, muscle force production, and fatigue. Specifically, these experiments tested two hypotheses: 1) whether an acute DOX exposure, at concentrations found in serum of cancer patients under chemotherapy, compromises RyR1 regular activity, and 2) whether pharmacological prevention of DOX-induced Ca^2+^ leak, using a novel class of triazole molecules (a.k.a. MP compounds) designed and synthesized for rescuing the FKBP-RyR1 interaction interface (Aizpurua *et al*., 2021; Passannante *et al*., 2023), can lessen the EC coupling dysfunction, thereby improving muscle force production and reducing DISM.

Our findings provide insights into potential therapeutic strategies that effectively alleviate DISM and that translate to positive downstream effects on muscle force production ex vivo. This discovery may hold the key to improving the well-being of cancer patients, potentially enhancing their adherence to chemotherapy.

## Materials and Methods

### Ethical approval

Protocols on the usage, care, and killing of animals were approved by the Institutional Animal Care and Use Committee of Rush University (IACUC). They were consistent with its ethical standards, as well as in compliance with the guidelines of the Animal Welfare Act.

### Animals

C57BL/6 wild-type (WT) mice, aged 15 to 20 weeks, were obtained from The Jackson Laboratories (Bar Harbor, ME). Animals were euthanized using CO_2_ inhalation followed by cervical dislocation. The *flexor digitorum brevis* (FDB) muscle was then carefully removed and immobilized in a custom 3D-printed chamber (Manno *et al*., 2017*a*) for subsequent Ca^2+^ or force measurements.

### Solutions

**Normal Tyrode** (mM): 140 NaCl, 5 KCl, 2.5 CaCl_2_, 2 MgCl_2_, 10 HEPES, 10 glucose, pH was adjusted to 7.2 with NaOH. **Chemical agents** (mM): 3.44 Doxorubicin hydrochloride (Cayman Chemical, Ann Arbor, Michigan, USA) was prepared in DMSO. 10 MP-001 and MP-034 (Miramoon Pharma S.L., San Sebastian, Spain) were dissolved in water and neutralized to pH 6.8 using NaOH.

**Single-channel recordings solutions** (mM): The cis-chamber contained 114 Tris, 250 HEPES, 0.5 EGTA, 5 total ATP, 1 free Mg, 0.1 free Ca^2+^, and pH 7.2. The trans-chamber contained 200 Cs-HEPES, 1 Ca^2+^, and pH 7.4. These solutions minimize/eliminate currents carried by these other non-RyR1 channels that may be present in the heavy SR vesicles.

**Calcium indicator** (mM): 5 Cal Red R525/650-AM (AAT Bioquest, USA) was prepared in DMSO. The dye was loaded by exposing muscles to a 10 µM solution of its AM derivative for 15 minutes at room temperature. Muscles were then washed with fresh Tyrode solution and allowed to rest for 15 minutes before measurements to enable de-esterification. **ROS indicator** (mM): 10 ROS Brite 670 (AAT Bioquest, USA) was prepared in DMSO and loaded for 20 minutes at 37°C at a final concentration of 10 µM.

Note: while the first muscle was used to perform the indicated DOX measurements, the contralateral muscle was preincubated with MP-001 for at least 2 hours.

### Single-channel recordings

Heavy SR microsomes were isolated from the gastrocnemius of rabbits. Single-channel recordings of RyR1 integrated into planar lipid bilayers were performed following a previously established method (Carvajal *et al*., 2024). In brief, the experimental setup consisted of two chambers, known as the cis and trans chambers. The cis chamber represented the cytosolic side and was held at virtual ground with the reference electrode. The trans chamber represented the luminal side and contained the voltage command electrode. SR microsomes were added to the cis-chamber and vigorously stirred before 50–300 μL of 4 M cesium methanesulfonate (Cs-CHOS) was added to promote microsome fusion. After microsome fusion, the cis-solution was exchanged for the cytosolic recording solution.

### Resting and transient cytosolic calcium concentration

The cytosolic Ca^2+^ concentration was calculated from the ratio 𝑅(𝑥, 𝑦, 𝑡) of fluorescence images 𝐹_1_ and 𝐹_2_ of Cal Red R525/650 excited with an Argon laser at 488 nm and collected between 500 and 575 nm (𝐹_1_) or between 600 and 725 nm (𝐹_2_), by the equation:

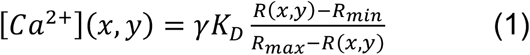

Here, γ is the ratio 𝐹_1_ of Ca^2+^-free dye /𝐹_1_ Ca^2+^-saturated dye. 𝑅_𝑚𝑖𝑛_ (0.01), 𝑅_𝑚𝑎𝑥_ (16), and γ𝐾_𝐷_ (3.3 μM) were determined by calibrations *in situ*. The resting values reported are averages over the area of cytosol spanned by the image.

### Calcium release flux

A typical line-scan image 𝐹(𝑥, 𝑡) and its components are shown in Fig. 4*A*. A time-dependent fluorescence ratio *R*(*t*) was calculated by averaging over the *x* coordinate for the extent of the image. Cytosolic Ca^2+^ concentration [Ca^2+^]_cyto_(*t*) was calculated by

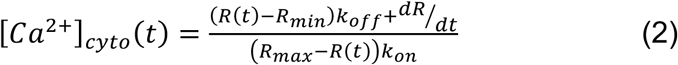

𝑅_𝑚𝑎𝑥_ and 𝑅_𝑚𝑖𝑛_ are the fluorescence intensity ratios at zero and saturating [Ca^2+^]. The Ca^2+^ release flux 𝑃(𝑡) was calculated from [Ca^2+^](*t*) using a simplified version of the “removal” method described previously (Melzer *et al*., 1984; Schuhmeier & Melzer, 2004), which involves assigning parameter values in a model of the removal of released Ca^2+^, so that the simulation fits the observed decay of cytosolic [Ca^2+^] simultaneously for several evoked field stimulation transients. For this procedure, we used parameters from various Ca^2+^ buffers and transport processes previously obtained from mammalian fast-twitch muscle fibers, as described by Baylor & Hollingworth, (2012).

Flux quantities were evaluated at specific times using a pair-pulse protocol for comparisons between control and DOX-treated muscles. The measured value was the early peak flux, which is also the absolute maximum, (*ṗ*_𝑡_). Residual fractions were calculated as the ratio between *ṗ*_2_/*ṗ*_1_ at 10 ms of interpulse interval (Jong *et al*., 1995).

### Contractile function

As a measure of whole muscle force production, we recorded tension generated by the FDB muscle ex vivo, as previously described (Manno *et al*., 2013*a*). Briefly, the three middle heads of the excised FDB muscles were tied/suspended between an ergometer motor (Aurora Scientific Model 407) and a fixed post, where the Achilles tendon was attached. The load cell was adjusted using a micromanipulator to stretch the muscle to a resting length between 0.9 and 1.1 cm. The cross-sectional area (CSA) was calculated by dividing the muscle volume by its length. The muscle volume was determined by dividing the muscle mass (in grams) by the density (assumed to be 1.06 g/cm³) (Engelke *et al*., 2018; Monte & Franchi, 2023).

FDB contraction was then triggered via electrical stimulation using two platinum electrodes placed in parallel to the muscle, and the resulting force was acquired and analyzed using pClamp software (version 10; Molecular Devices, San Jose, CA, USA). We employed the following stimulation protocols:

- For **force-frequency measurements** (weakness), muscles at optimal sarcomere length were stimulated with tetanic stimulations at various frequencies (1, 5, 10, 20, 40, 80, 100, 120, & 150 Hz) with 2 minutes recovery period between episodes to minimize fatigue.
- For **fatigue measurements**, tetanic stimulations (80 Hz for 350 ms) were applied every 2 s for 5 minutes.

All experiments, ranging from single-channel recordings to measurements of muscle force and fatigue, were conducted at room temperature (20-22°C).

### Statistics and presentation of replicates

The measurements compared in the present study are typically quantitative features of *N* individuals (mice) with *n*_i_ replications (muscles) for individual mouse *i*. Statistical significance of the difference between variables was determined using hierarchical (or nested) analysis (Sikkel *et al*., 2017; Lazic *et al*., 2018). Means were calculated conventionally, averaging over all measurements, for example:

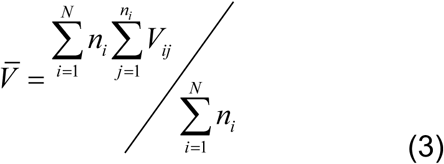

To assess the statistical significance of differences, the effective number of cases was calculated. This calculation accounts for the grouping or clustering of replicates from the same individual, thus avoiding pseudoreplication (Lazic *et al*., 2018). The effective number is smaller than the actual number of replicate measures, which results in a higher SEM and an increased *p*-value for the null hypothesis of no difference. Therefore, the nested analysis helped prevent overestimation of significance by utilizing the information available from the replicates.

## Results

### Doxorubicin interferes with muscle contraction

Weakness and fatigue are among the side effects caused by doxorubicin (DOX) in cancer patients (Neidhart *et al*., 1984). In this work, we explored the side effects of DOX and tested a strategy to mitigate them. To gain a better understanding of Ca^2+^ management in mouse muscles treated with this chemotherapeutic agent at serum levels similar to those found in cancer patients, we combined established techniques for dynamic measurements of cytosolic Ca^2+^ transients and the underlying Ca^2+^ release flux. We also studied the extent of recovery from inactivation of Ca^2+^ release in response to single action potentials, by using a double pulse technique in which the effect of a conditioning stimulation is determined by the change in Ca^2+^ release elicited by a second stimulation, applied at different time intervals (Jong *et al*., 1995; Caputo *et al*., 2004). Finally, we assessed the rate of reactive oxygen species to determine their importance in the unbalanced Ca^2+^ homeostasis induced by DOX.

#### DOX induces FDB muscle weakness, and MP-001 prevents it

To investigate the acute impact of DOX exposure and the beneficial effect of MP-001 on isometric force production, ex vivo FDB muscles were incubated for 2 h in Tyrode solution containing 10 µM DOX, a dose corresponding to serum levels in patients and animals undergoing chemotherapy (Morris *et al*., 1989; Swenson *et al*., 2003; Sakai-Kato *et al*., 2012). The contralateral muscle was pre-incubated with 10 µM MP-001. Figure 1***A*** depicts an example of representative absolute force superimposed signals (in grams), acquired at increasing electrical stimulation frequencies (trains of 350 ms duration, from 1 to 120 Hz). Signals were acquired at 1 and 2 hours of incubation with DOX alone (*a1*). After 2 hours of preincubation with MP-001, DOX was added, and samples were measured at 1 and 2 hours (*a2*). In this example, the entire set of signals decreased force production by 30% after 2 hours, compared to a 10% decrease when preincubated with MP-001. Figure 1***B*** shows a summary of the specific force acquired at different frequencies and normalized to the cross-sectional area (CSA). The maximum force produced at the last two frequencies (100 and 120 Hz) decreased by ∼8% and 15% after 1 and 2 hours, respectively, after applying DOX. This continuous impairment in force production was improved when the muscles were preincubated with MP-001, resulting in a 4% and 9% decrease in force, as shown in (*b1*), and is summarized at 100 Hz in (*b2)*; \**P* < 0.05; N = 4. We did not find significant differences in the kinetics of these curves, as shown in Fig. S1. However, the maximum contraction values reached at all frequencies were significantly higher when preincubated with MP-001 compared to the control + DOX, as shown in *b1* (\**P* < 0.05), and summarized in Table 1.

**Figure 1.**
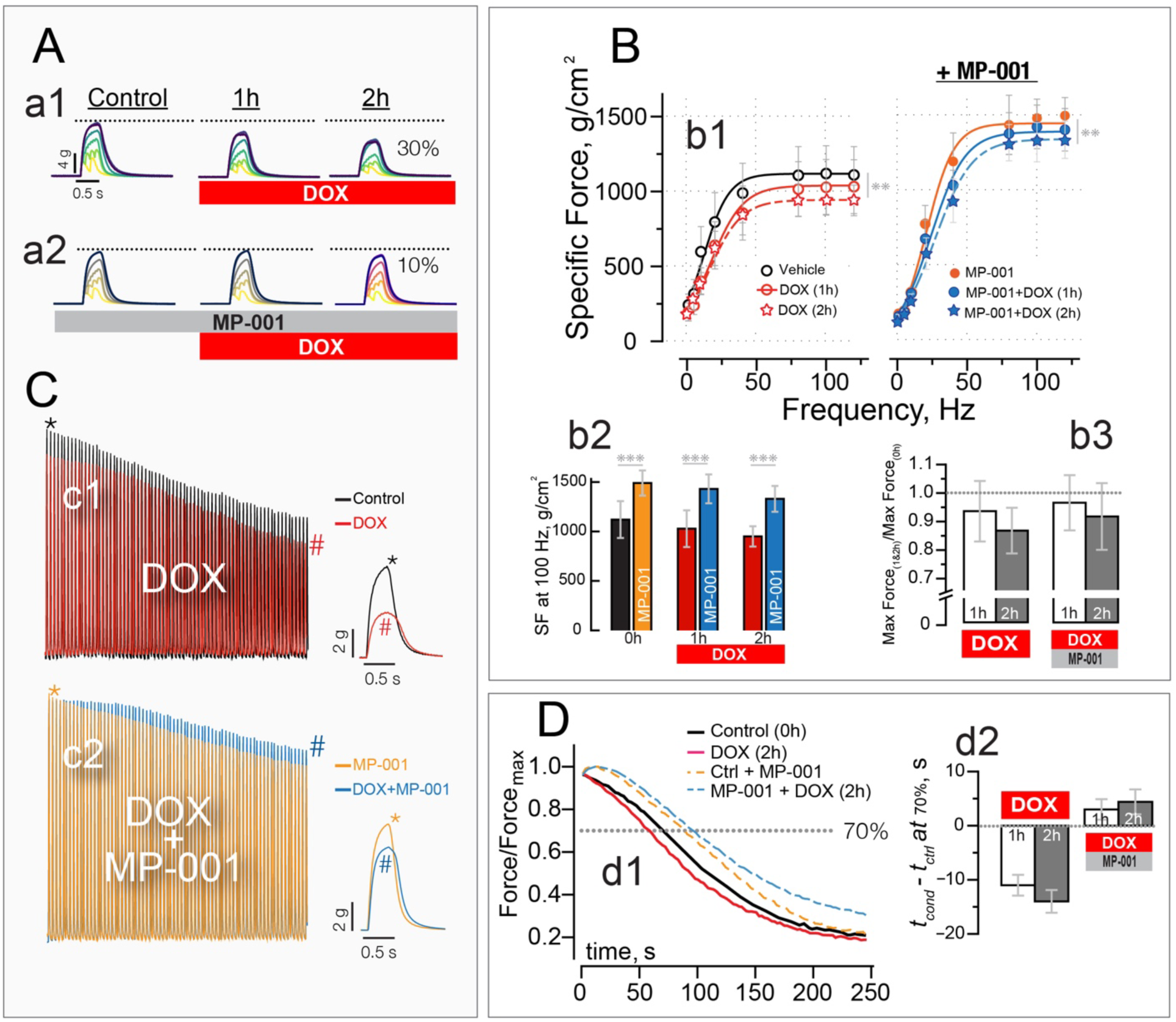
DOX induces weakness and fatigue, and MP-001 prevents these effects. <u>Muscle force</u> <u>production</u>: Panel ***A***, ex vivo FDB muscle contractile force responses to trains of 350 ms duration stimuli at frequencies from 5 to 120 Hz, with DOX (10 µM, in ***a1***), and after pre-incubation for a minimum of two hours with the reshaper MP-001 (10 µM, ***a2***). Measurements were performed after 1 and 2 hours of the addition of DOX. For a summary of maximum peak values obtained at different frequencies, see panel ***B*** (***b1***), where sigmoidal functions were plotted using the average of the fitting parameters in an N = 4 muscles per condition; \*\**p* < 0.01 in two-level hierarchical analysis (versus the comparative data set). In ***b2***, we compare the maximum specific force generated at 100 Hz, and in ***b3***, we normalize them with the control values, between DOX and muscles pre-incubated with MP-001. <u>Fatigue</u>: ***C,*** A fatigue protocol of 80 Hz, 350 ms, every 2 seconds for 5 minutes was applied on FDB muscles. Typical superimposed signals of total force acquisition in (***c1 & c2***), where black/orange traces represent control and control + MP-001, respectively. Red/blue traces represent DOX and DOX + MP-001 present, respectively, after two hours. ***D*** (***d1***) Summary of normalized signals of maximum force decay values, as shown in ***C***. Panel ***d2*** shows the average time it takes to fatigue the muscles by 30% of the initial force production. All data are presented as mean ± SD; \*\**p* < 0.01 in two-level hierarchical analysis (versus the comparative data set). N=4.

**Table 1.** Descriptive statistics for force and fatigue parameters.

| | | | *Specific Force<sup>#</sup>*<br>*(g/cm<sup>2</sup>)* | *Time to reach*<br>*70% of force<sup>@</sup> (s)* | *% of fatigue*<br>*reached<sup>\$</sup>* |
| --- | --- | --- | --- | --- | --- |
| DOX (10 μM) | -MP-001 | 0h | 1112.86 ± 188 | 72 ± 7 | 18.8 ± 4 |
| | | 1h | 1025.33 ± 182 | 61 ± 7 | 21.7 ± 4 |
| | | 2h | *944.4 ± 102 | *58 ± 8 | 21.9 ± 5 |
| | +MP-001 | 0h | *1480.46 ± 190 | 88 ± 7 | 28.7 ± 5 |
| | | 1h | *1410.13 ± 142 | *91 ± 7 | 27.6 ± 4 |
| | | 2h | *1333.42 ± 225 | *92.4 ± 9 | 24.7 ± 3 |

Values are means ± SEM. \* $P$ < 0.05 in two-level hierarchical analysis.
Values were compared to those of the control without MP-001.

#### DOX induces fatigue, and MP-001 prevents it

Figure 1*C* shows representative traces acquired using a fatigue protocol consisting of 350 ms tetanic stimulations at 80 Hz, repeated every 2 seconds for 5 minutes. Panels *c1* and *c2* depict typical whole traces with DOX and DOX + MP-001, respectively. In this set of experiments, a muscle was deemed fatigued when its contraction force decreased by 30%, as indicated by the dotted line in panel *d1* and summarized in panel *d2*, where the time it takes to reach 70% of the initial contraction force becomes shorter when muscles are treated with DOX. MP-001 delays the time it takes for muscles to fatigue in all cases.

### Doxorubicin directly activates RyR1

#### Quantifying Single RyR1 Function

To better understand the complex actions of DOX and the normalizing influence of the FKBP12 reshapers on muscle force production, shown in the previous section, we decided to start by measuring the actions of these drugs on single basal RyR1 channel function at 100 nM free cytosolic (cis) Ca^2+^ present. Figure 2A shows multi-RyR1 sample recordings (upward deflections indicating openings) at 0 and 120 minutes after DOX and DOX + MP-001 application. The initial rise in single RyR1 open probability (Po) after DOX application is plotted in Figure 2B (black line). The DOX action on Po did not occur if MP-001 was present (red line). The single RyR1 open probability (Po), frequency of openings longer than 6 ms (FREQ>6ms) and mean open time (MOT) results (mean ± SEM; n=8) in the absence (black circles) and presence of an FKBP reshaper (MP-001 red, MP-034 blue circles) without DOX present (control) and two hours after applying DOX are shown in Figure 2*C, D* and *E*. MP-001 & MP-034 by themselves did not affect single RyR1 Po, FREQ>6ms, and MOT. DOX evoked an initial increase and later decrease (not shown) in single RyR1 Po, FREQ>6ms, and MOT. The Po remained significantly elevated (compared to control) at the 2-hour mark (panel ***A***). The subsequent addition of MP-001 or MP-034 normalized the DOX-evoked changes in single RyR1 Po, FREQ > 6ms, and MOT. In most cases, MP-001 was more effective than MP-034 in DOX’s action on single RyR1 function.

**Figure 2.**
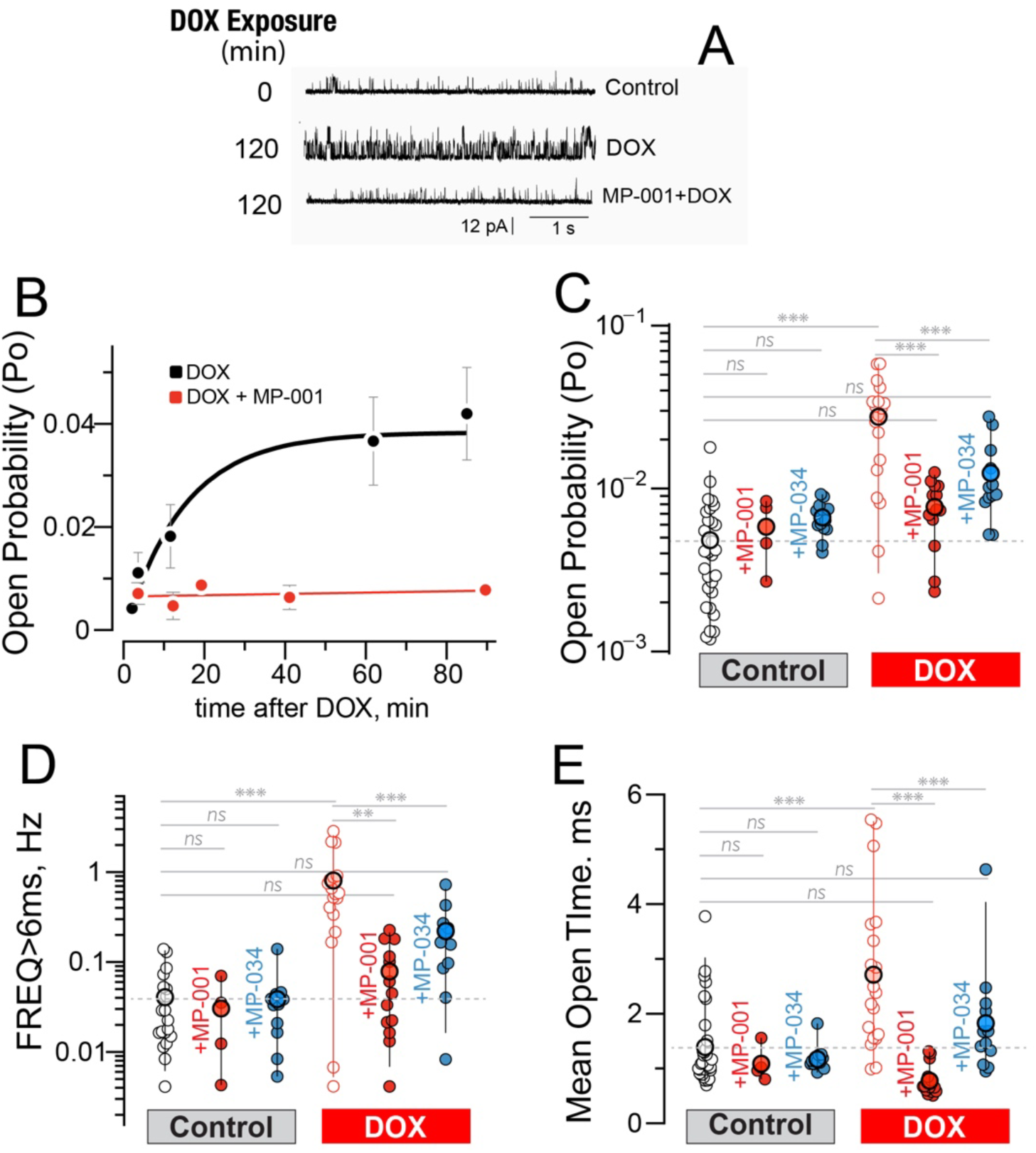
RyR1 activity is increased by DOX and prevented by MP-001/MP-034. Panel ***A***, example single-channel recordings acquired at the times indicated after DOX and MP-001+ DOX exposure. Open events are shown as upward deflections. Single RyR1 activity was acquired with 100 nM free cytosolic (cis) Ca present. ***B***, DOX (10 µM) activated basal single RyR1 Po progressively over time after application (black line); red line shows that activity remains constant when treated with MP-001 together with DOX. ***C***, ***D***, and ***E***, DOX significantly increased the basal RyR1 Po, FREQ>6ms, and mean open times (MOT) after 3 minutes, and this was sustained for 2 hours. The subsequent application of MP-001 or MP-034 was acquired 2 hours after application of either drug together with DOX. Small symbols represent single experiments, bigger circles are mean values, and lines represent a 95% confidence interval for obtaining a point within such a range. \*\**p* < 0.01, \*\*\**p* < 0.001 (ANOVA, versus the comparative data set); ns, not significant.

### Calcium-dependent inactivation of RyR1 is affected by DOX

#### Calcium-dependent inactivation (CDI) was studied with field stimulation

The CDI index, measured as the residual fraction (RF) of Ca^2+^ release remaining after the second electrical stimulation, changes over time. We used the whole FDB muscle stretched in our 3D-printed chambers (Figure 3A). Muscles were loaded with Cal Red-AM and field-stimulated with platinum electrodes aligned parallel to the fibers. To better resolve the kinetics of this event, we focused solely on the upward emission at 525 nm from this ratiometric dye. Panel *a1* illustrates fluorescence traces associated with field stimulation using a paired-pulses protocol with intervals ranging from 10 ms to 200 ms apart. CDI was calculated as the ratio of the peak fluorescence signal at the second peak to that at the first peak, F_2_/F_1_ (*a2*), and summarized for all the time intervals in *a3*. Figure S2*A* summarizes the signals obtained at each stimulation frequency and under the three different conditions: Control (*a1*), DOX (*a2*), and DOX + MP-001 (*a3*). Interestingly, DOX progressively decreases the CDI in these muscle fibers, with the most significant change occurring at 10 ms. In panel 3***B***, we summarize the ratio that reflects the degree of inactivation occurring at 10 ms interval between pulses. The curves in panel 3***C*** represent the kinetic changes in fractional peak values for all stimulation frequencies, as shown in Fig. S2*A*. These curves were generated by plotting an exponential-saturating function (summary of parameters in Table 2) with the average parameters fitted to each data set (summary of values in Table 3), at two hours control (*N* = 5, *n =* 27), after application of DOX (*N* = 5, *n =* 14) and DOX + MP-001 (*N* = 4, *n =* 8). Shadow regions represent the non-inactivating component projected at the 10 ms intersection. Values increased from a 14.2% fractional peak in control to a 61.4% after 2h in DOX. Fraction peak values increased only by 25% when fibers were pretreated with MP-001 (Table 2).

**Figure 3.**
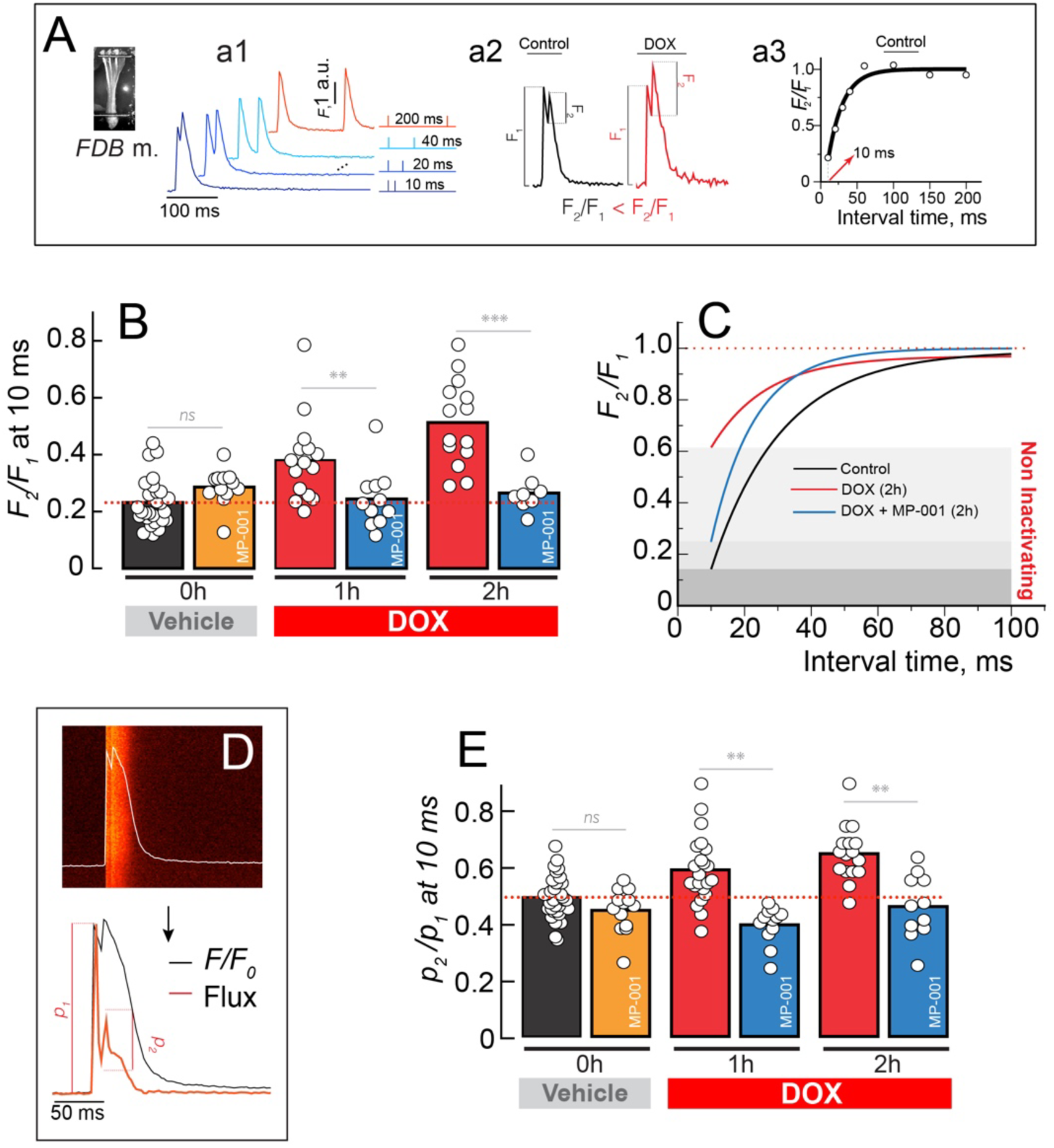
Recovery from inactivation of Ca release after paired-pulse electrical stimulation. ***A, a1***, Signals were elicited by two action potentials separated at interpulse intervals of 10-200 ms, in FDB muscles fixed in a 3D-printed chamber (inset). In ***a2,*** it is shown that the fractional peak value (**FP**) of Ca-dependent fluorescence is the ratio of the second peak to the first peak (*F_2_/F_1_*), and an example is plotted for each time interval in ***a3***. In ***B,*** we plot the FP values at 10 ms interval for each condition. Values are significantly different between control and condition at 1 & 2 hours, \*\**P < 0*.01 & \*\*\**P < 0*.001 in two-level hierarchical analysis. We also plot the FP values vs each interpulse interval. The average of the fitting parameters for each condition is used to plot the exponential function shown in ***C***. The shadow areas represent the non-inactivating components at a 10 ms interval, after 2 hours, for control, DOX, and DOX + MP-001. In ***D,*** flux values were calculated from the fluorescence signal as explained in the methods. Panel ***E*** depicts the residual fraction (**RF**) calculated for signals under the different conditions, and represented as the ratio of the 2^nd^ peak of Ca^2+^ flux to the value at the 1^st^ peak (*p_2_/p_1_*), similar to ***B***. Values are mean, *ns* represents not significant, and \*\**p*<0.01 in two-level hierarchical analysis.

**Table 2.**
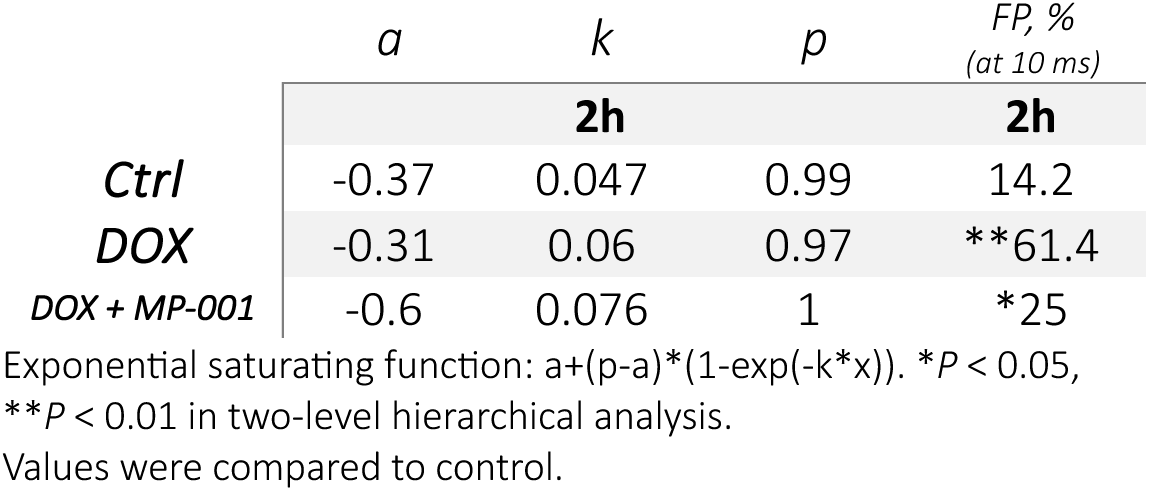
Parameters of Residual Fraction Recovery of Ca^2+^ Release after Paired Pulses.

|  | *a* | *k* | *p* | *FP, %*<br>(at 10 ms) |
| --- | --- | --- | --- | --- |
|  | **2h** |  |  | **2h** |
| *Ctrl* | -0.37 | 0.047 | 0.99 | 14.2 |
| *DOX* | -0.31 | 0.06 | 0.97 | **\*\*61.4** |
| *DOX + MP-001* | -0.6 | 0.076 | 1 | **\*25** |

Exponential saturating function: $a + (p - a) * (1 - \exp(-k * x))$ . \* $P < 0.05$ , \*\* $P < 0.01$ in two-level hierarchical analysis.
Values were compared to control.

**Table 3.**
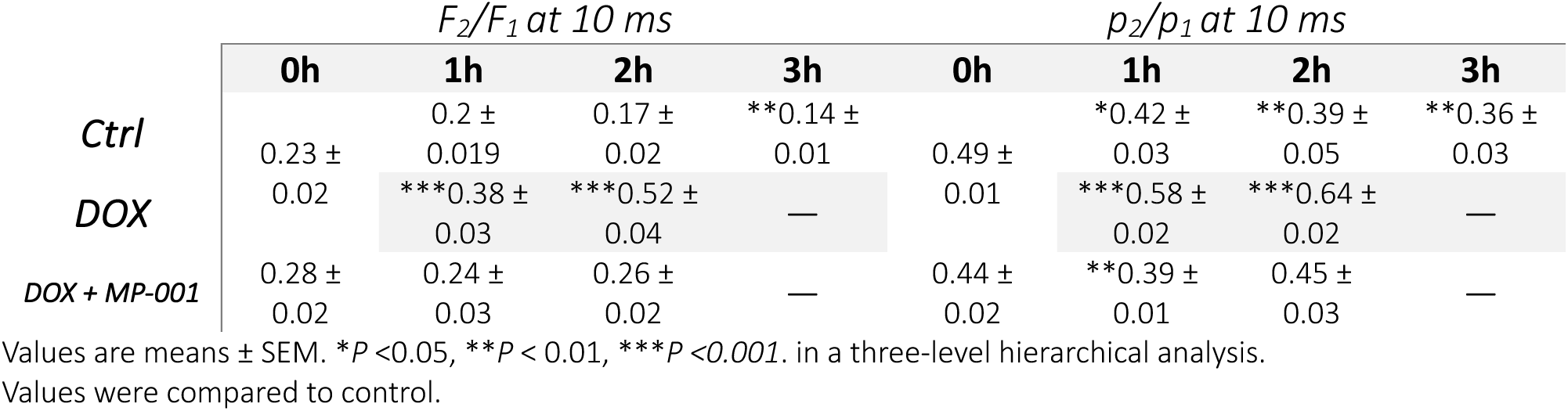
Descriptive Statistics for the Recovery of Ca^2+^ Release after an Action Potential.

#### Calcium release flux was affected by DOX

To reduce variability from fluorescence signals, we improved the quantification of CDI by calculating the Ca^2+^ release flux, which represents the actual rate of Ca^2+^ movement, using a simplified removal method (Fig. 3D). From the recordings with the shortest interval between action potentials, specifically, the 10 ms gap between pulses as shown in Fig. 3a2, the peak values of the flux signals were used to calculate the residual fraction (RF), defined as RF = *p_2_/p_1_*, at the 10 ms interval. These values were considered as the non-inactivating component of Ca^2+^ release. When we calculated these values under control conditions, both with and without MP-001, we found that RF did not differ significantly. Notably, under control conditions, the fraction of Ca^2+^ release decreased over time, showing a 25% drop after three hours, which suggests increased Ca-dependent inactivation (Fig. S3***C***). Conversely, DOX gradually increased Ca^2+^ release at the peak of the second depolarization, reaching a 30% increase after two hours (Fig. 3E). Values obtained after preincubation with MP-001 decreased after one hour, but were not significantly different from control throughout time.

### Doxorubicin affects calcium homeostasis

After analyzing the results of single RyR1 channels and Ca-dependent inactivation, we rationalized that DOX would likely disrupt normal intracellular Ca^2+^ homeostasis. To verify this conclusion, we recorded basal steady state Ca^2+^ and Ca^2+^ transients elicited by prolonged high-frequency stimulation-induced action potentials after loading the muscle with Cal Red R525/650. Figure 4A shows the representative signals from this ratiometric Ca^2+^ indicator; the emission signal increases at 525 nm (*a1*) and decreases at 650 nm (*a2*) when excited at 488 nm. The ratio of F_1_/F_2_ is shown in *a3*. Typical Ca-calibrated signals, derived from parameters in the Methods section, are displayed in Figure 4B under the three conditions: vehicle (DMSO), DOX, and DOX + MP-001. These signals consisted of Ca^2+^ transient signals triggered by 120 Hz trains lasting 350 ms, repeated every hour.

#### Basal cytosolic Ca^2+^ concentration is affected by DOX and recovered by MP-001

The calibration of Cal Red allowed us to follow the changes in basal cytosolic Ca^2+^ concentration after the application of DOX and DOX + MP-001. Figure 4C illustrates these concentrations at 1 and 2 hours, while Figure S3***A*** displays the percentage changes relative to the initial [Ca^2+^]_cyto_ value (represented as 0h). We observed an increase from 105 ± 3.7 nM at basal conditions to a maximum mean concentration of 153 ± 4.4 nM, a 31% increase after 2 hours (see Table 4). We must note that, although we corrected for the diffusion of DOX into the fibers, these values may be underestimated due to the slight increase in DOX-related fluorescence at 650 nm, which overlaps with the bandwidth of the second channel (*a2*). Interestingly, basal [Ca^2+^]_cyto_ remained constant during the 2h pre-incubation with MP-001, reaching only 115 ± 5.9 nM or 0.76. (Table 4).

**Table 4.**
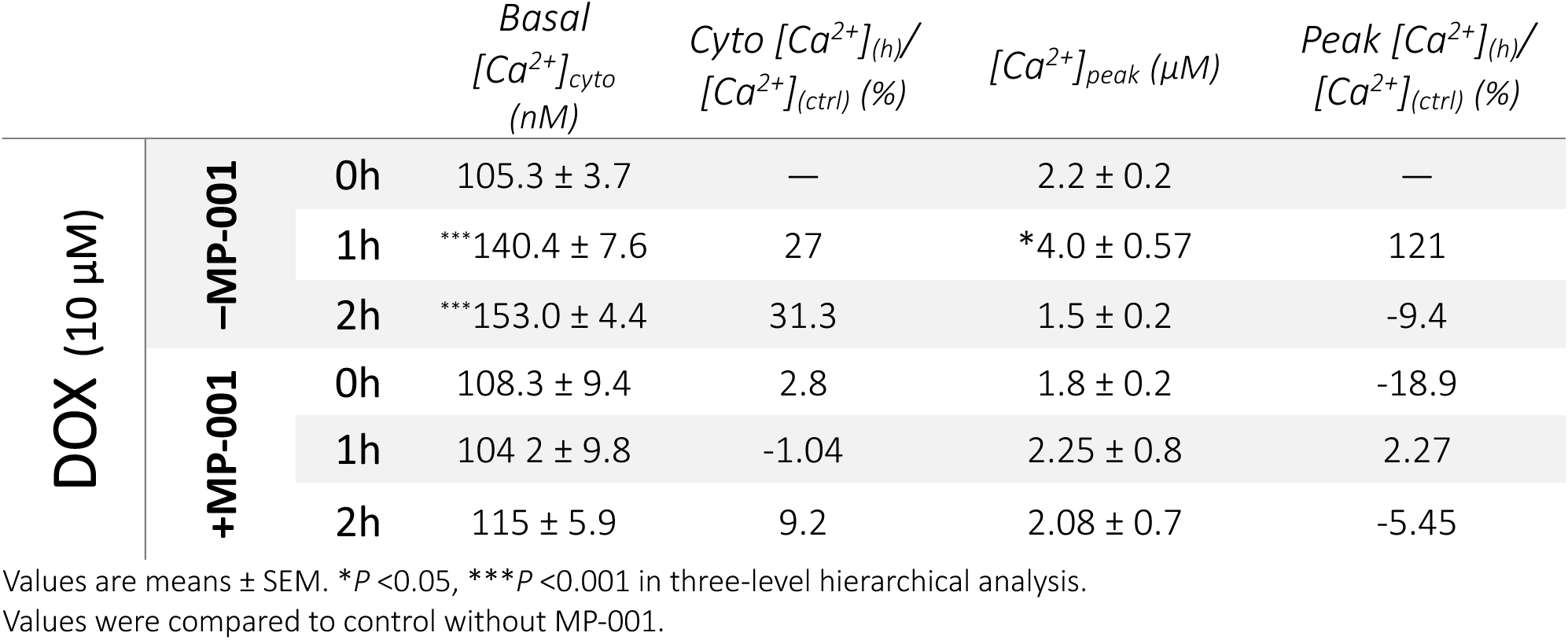
Descriptive statistics for basal and dynamic calcium changes.

|  |  |  | Basal<br>[Ca<sup>2+</sup>]<sub>cyto</sub><br>(nM) | Cyto [Ca<sup>2+</sup>]<sub>(h)</sub>/<br>[Ca<sup>2+</sup>]<sub>(ctrl)</sub> (%) | [Ca<sup>2+</sup>]<sub>peak</sub> (μM) | Peak [Ca<sup>2+</sup>]<sub>(h)</sub>/<br>[Ca<sup>2+</sup>]<sub>(ctrl)</sub> (%) |
| --- | --- | --- | --- | --- | --- | --- |
| DOX (10 μM) | -MP-001 | 0h | 105.3 ± 3.7 | — | 2.2 ± 0.2 | — |
|  |  | 1h | ***140.4 ± 7.6 | 27 | *4.0 ± 0.57 | 121 |
|  |  | 2h | ***153.0 ± 4.4 | 31.3 | 1.5 ± 0.2 | -9.4 |
|  | +MP-001 | 0h | 108.3 ± 9.4 | 2.8 | 1.8 ± 0.2 | -18.9 |
|  |  | 1h | 104.2 ± 9.8 | -1.04 | 2.25 ± 0.8 | 2.27 |
|  |  | 2h | 115 ± 5.9 | 9.2 | 2.08 ± 0.7 | -5.45 |

Values are means ± SEM. \**P* < 0.05, \*\*\**P* < 0.001 in three-level hierarchical analysis. Values were compared to control without MP-001.

#### Action potential-initiated Ca^2+^ release is affected by DOX and recovered by MP-001

We also quantified changes in the peak Ca^2+^ release, as shown in Fig. 4D, and summarized its percentage relative to the control value (2.2 ± 0.2 µM), shown in Fig. S3***B***. Interestingly, DOX caused a sustained increase in Ca^2+^ signal amplitude, which significantly peaked within the first hour of application at 4.0 ± 0.57 µM, representing a 121%. Then, after 2 hours, the amplitude decreased to values below the control level (1.5 ± 0.2 µM), indicating a 9.4% reduction. A consistent feature obtained was the lack of changes when MP-001 was used in conjunction with DOX. Further rate-dependent changes are depicted in Table 4.

**Figure 4.**
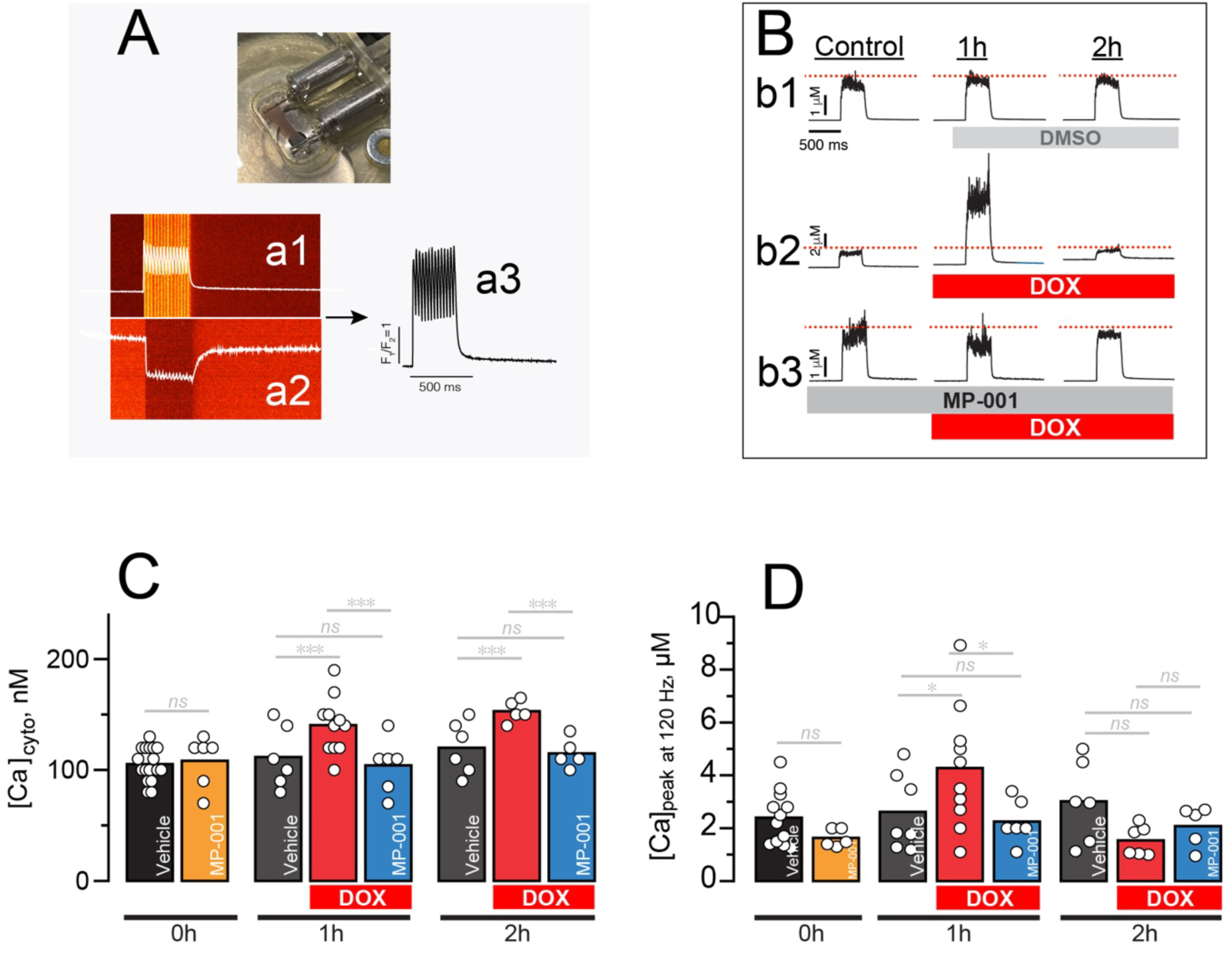
DOX increases & MP-001 restores peak and basal cytosolic Ca^2+^. Panel ***A***, Cell-wide Ca^2+^ transients measured by linescan imaging in stretched muscle fibers mounted in a 3D-printed chamber, and using the ratiometric Ca indicator Cal Red. In panel ***a3***, the ratio (*F_a1_/F_a2_*) signal between the two emission channels is represented. ***B,*** typical Ca^2+^ transients elicited by electrical stimulation. Panel ***b1*** shows representative data, where DMSO was used as the vehicle, displaying minimal signal change, with an approximately 10% decrease in release after two hours. Panel ***b2*** shows representative traces after exposing muscles to DOX, and in panel ***b3***, after preincubating with MP-001. ***C*** & ***D*** are summaries of basal cytosolic and peak [Ca^2+^] before and after DOX and DOX + MP-001 application. Values are mean, *ns* represents not significant, \**p*<0.05, & \*\*\**p<0.00*1. in a two-level hierarchical analysis. N=4.

### Doxorubicin produces reactive oxygen species in response to elevated cytosolic Ca^2+^ levels

Mitochondria are a significant source of DOX-induced reactive oxygen species (ROS) production (Min *et al*., 2015). Therefore, we aim to determine whether ROS production, in addition to the direct action of DOX within the mitochondria, is also dependent on the increase in cytosolic Ca^2+^. This will also clarify the extent to which the increased [Ca^2+^]_cyto_ is derived from ROS oxidative effects on Ca^2+^ release channels, as previously suggested (Gilliam *et al*., 2013*b*).

Oxidative stress produced by DOX was measured by assessing ROS levels using ROS Brite 670. Figure 5***A*** *&* ***B*** show that ROS production becomes faster and reaches higher values (*b1 & b2*) when DOX is applied, compared to the normal ROS production observed under control conditions. Specifically, the maximum slope of ROS production under control conditions was 0.0146 min^-1^, whereas it increased to 0.0427 min^-1^ with DOX, approximately 3 times the control value. Based on the values observed in panel *B (b1)* and summarized in Table 5, we calculated that it would take 78 minutes to increase from 10% to 90% of its ROS-related fluorescence value. In contrast, when DOX was present, it would only take approximately 25 minutes to reach the same value. When we preincubated with MP-001, it took approximately 100 minutes to reach 90% from 10%, about 13% longer than the control group. Interestingly, when we used the reshaper molecule MP-034, which lacks the additional antioxidant properties of MP-001, the curve was approximately 4.4% steeper and faster. MP-034 reached 90% about 8.8 minutes earlier than MP-001, but was still 12.6 minutes slower than the control conditions.

**Figure 5.**
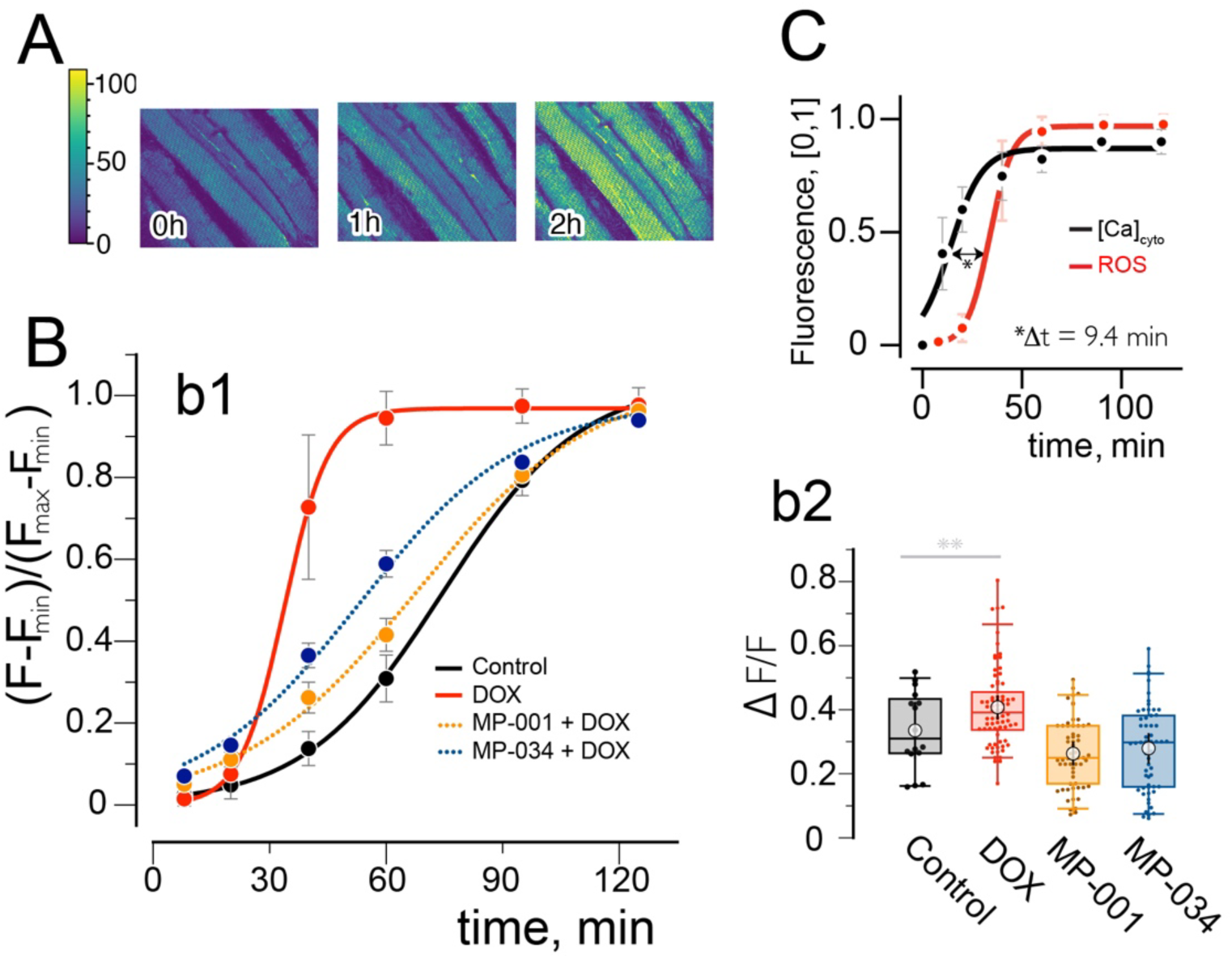
DOX-driven ROS production. ***A***, FDB fibers loaded with ROS-brite 670 show an increase in ROS-dependent fluorescence after applying DOX (10 µM). ***B***, DOX-driven ROS production in red, abruptly increased in comparison with the control (black). In addition to its reshaper effect, MP-001 (orange) has antioxidant activity. We used a non-antioxidant version (MP-034) to determine the importance of this effect (blue curve). The maximum amplitude of fluorescence, reached after 2 hours, is summarized in ***b2***. ***C***, Normalized fluorescence of ROS together with the dynamic increase in basal cytosolic Ca^2+^. Values are mean ± SD, \*\**p*<0.01 in three-level hierarchical analysis.

**Table 5.** Parameters of ROS production.

|  | *a* | *b* | *x<sub>0</sub>* | *Max Slope,*<br>*min<sup>-1</sup>* | *ROS, 10-90%*<br>*Time, min* |
| --- | --- | --- | --- | --- | --- |
| *Ctrl* | 1.04 ± 0.01 | 17.8 ± 0.48 | 74.6 ± 0.8 | 0.0146 | 78.15 |
| *DOX* | 0.97 ± 0.006 | **5.7 ± 0.25 | **33.8 ± 0.36 | *0.0427 | **24.96 |
| *DOX + MP-001* | 1.04 ± 0.04 | 22.6 ± 1.76 | 67.8 ± 2.8 | 0.0114 | *99.56 |
| *DOX + MP-034* | 0.98 ± 0.008 | 20.6 ± 1.5 | 53 ± 1.6 | 0.0119 | 90.79 |

Parameters used to fit the logistic function $a/(1+\exp(-(x-x_0)/b))$ .
Values are means ± SEM. \**P* < 0.05, \*\**P* < 0.01 in three-level hierarchical analysis. Values were compared to control.

Following the same rationale, we plotted the curve of ROS production alongside the increase in Ca^2+^ concentration after adding DOX, in panel *C.* We observed that the increase in ROS production was preceded by the release of Ca^2+^, which occurred approximately 10 minutes earlier. This suggests that the impact of DOX on RyR1 is more significant than merely controlling ROS production. Consequently, we proposed that skeletal muscle dysfunction following DOX exposure is a two-fold process resulting from i) a direct effect on the RyR1, likely destabilizing the interaction between RyR1-FKBP12, and ii) a direct toxic effect on mitochondria leading to increased ROS production. In this context, ROS can oxidize RyR1 channels, disrupting Ca^2+^ homeostasis and creating a vicious cycle of Ca^2+^ overload and oxidative stress that negatively affects muscle contraction.

## Discussion

The present study offers new insights into the acute effects of doxorubicin (DOX) on skeletal muscle function, with a particular focus on the mechanisms underlying Doxorubicin-Induced Skeletal Myotoxicity (DISM).

Our ex vivo experiments demonstrate that DOX, at clinically relevant concentrations, impairs both isometric force production and resistance to fatigue in isolated FDB muscles. These findings are consistent with previous studies on DOX-induced myopathy (Ertunc *et al*., 2009) and extend our understanding by dissecting the molecular events that contribute to contractile dysfunction. The reduction in force production following DOX treatment is attributed to mitochondrial respiratory inhibition and elevated ROS production (Sarvazyan, 1996; Xiong *et al*., 2006; Gilliam & Clair, 2011; Gilliam *et al*., 2013*a*). Nevertheless, it has been found that, even though antioxidants prevent DOX-induced mitochondrial ROS production and protect against skeletal muscle atrophy and contractile dysfunction, they do not completely rescue these defects, indicating that ROS scavenging alone is insufficient to prevent these side effects (Yang *et al*., 2014; Min *et al*., 2015; Tarpey *et al*., 2019; Pigg *et al*., 2025).

In this study, we found that, beyond DOX’s direct effects on mitochondrial function, prolonged elevation of cytosolic [Ca^2+^] significantly increases ROS production. High Ca^2+^ concentration in the microdomain between RyR1 and mitochondria (Marcucci *et al*., 2023) raises intramitochondrial [Ca^2+^], which alters the Krebs cycle and electron transport chain activity, and activates nitric oxide synthase (NOS), further promoting ROS generation (Gilliam *et al*., 2013*a*; Görlach *et al*., 2015; Feno *et al*., 2019). Thus, controlling this Ca^2+^ dysregulation is key to improving muscle performance during chemotherapy.

### Is the RyR1 affected by DOX?

In line with a chemical-caused couplonopathy, and because multiple proteins within the triad are allosterically and structurally connected, DOX can affect one or more couplon proteins, disrupting related Ca^2+^ release mechanisms and thus causing similar phenotypes. An example of this is the functional and structural effects of high doses of DOX on altered calsequestrin polymerization and SR Ca^2+^ store depletion (Subra *et al*., 2012). Although our study did not directly investigate calsequestrin dynamics, the long-term DOX-induced Ca^2+^ leak observed may also destabilize the calsequestrin network, further impairing SR Ca^2+^ buffering and directly affecting Ca^2+^ release (Manno *et al*., 2017*b*).

Our findings are consistent with previous studies, including our own, which show that persistent activation or destabilization of RyR1, whether caused by pharmacological agents, genetic ablation of stabilizing proteins, or oxidative modifications, leads to an increase in SR Ca^2+^ leak (Manno *et al*., 2013*c*, 2022; Canato *et al*., 2019). Specifically, studies have shown that DOX destabilizes RyR channels, affecting both single RyR2 channels (Hanna *et al*., 2014) and skeletal muscle vesicles carrying RyR1 (Zorzato *et al*., 1985*b*). A key finding in this work is the increased activity of RyR1 after DOX exposure. This is evidenced by a higher open probability (Po) and longer mean open times (MOT) observed in our single-channel recordings, along with a reduction in Ca-dependent inactivation observed at the muscle fiber level. This abnormal RyR1 activation likely supports our hypothesis that an enhanced SR Ca^2+^ leak disrupts Ca^2+^ homeostasis, ultimately leading to muscle weakness.

Multiple studies have highlighted that calcium-dependent inactivation (CDI) and the repriming of the RyR1 channel are tightly regulated by both the amplitude and kinetics of Ca^2+^ transients, as well as by the intrinsic properties of the RyR1 channel and its associated couplon proteins (Jong *et al*., 1995; Caputo *et al*., 2004). For example, it has been demonstrated that CDI is modulated by the local [Ca^2+^] near the RyR1 channel pore, and that alterations in SR Ca^2+^ buffering, such as those caused by Casq1 depolymerization, can significantly impact the inactivation process. Our data indicate that DOX increases basal [Ca^2+^]_cyto_ and disrupts SR Ca^2+^ homeostasis. This aligns with previous findings and suggests that DOX-induced SR Ca^2+^ leak directly impairs the mechanisms of CDI.

Furthermore, the increased repriming of RyR1 caused by DOX, as shown by the reduced inactivation fraction in paired-pulse protocols, suggests that DOX not only increases SR Ca^2+^ leak but also disrupts the normal feedback mechanism that limits Ca^2+^ release during repetitive stimulation (Schneider & Simon, 1988). This effect is especially noticeable at short interpulse intervals, where the residual fraction of Ca^2+^ release (F_2_/F_1_) is higher in the presence of DOX. This indicates an enhanced repriming process or, in other words, an increased ability of the inactivated RyR1 channels to return to a state of readiness for subsequent activation. These findings support our hypothesis that DOX weakens the protective role of CDI against excessive Ca^2+^ release during sustained activity, as previously described (Jong *et al*., 1995; Manno *et al*., 2013*b*).

The dual effect of increased leak and decreased SR Ca^2+^ levels exacerbates cytosolic Ca^2+^ overload by activating store-operated Ca^2+^ entry (Arora *et al*., 2021; Nemoto *et al*., 2022). In this way, our findings support the notion that an increase in basal cytosolic Ca^2+^, along with altered kinetics of Ca^2+^ transients following DOX treatment, likely contributes to increased susceptibility to fatigue and weakness. This occurs as muscle fibers lose their ability to self-limit Ca^2+^ release during repetitive activation (Schneider & Simon, 1988). Furthermore, elevated levels of free cytosolic Ca^2+^ activate calpains and, over time, promote caspase-3 activation, which can be detrimental to normal muscle contraction (Min *et al*., 2015).

In addition to the direct consequences of inducing a prolonged high [Ca^2+^]_cyto_, it has been thoroughly described that DOX induces other metabolic dysregulations (de Lima Junior *et al*., 2016), consistent with those observed by our group in models of Ca^2+^ dysregulation and calpain activation (Tammineni *et al*., 2020, 2025*a*, 2025*b*).

### DOX likely affects RyR1 by altering the interaction between RyR1 and FKBP12

It has been previously demonstrated that in specific pathologies, oxidized RyR1 channels result in the loss of RyR1-FKBP12 interaction; thus, restoring it can reduce Ca^2+^ leak, thereby decreasing ROS production and oxidative stress (Waning *et al*., 2015). In the process of rescuing the Ca^2+^ dysregulation induced by DOX, we tested MP-001, a drug designed and synthesized to target and rescue the RyR1-FKBP12 protein-protein interaction (Aizpurua *et al*., 2021). We found that this drug can mitigate many of the deleterious effects of DOX on muscle function. MP-001 preserved force production, delayed the onset of fatigue, and normalized both single RyR1 activity and cytosolic Ca^2+^ levels. Therefore, this protective effect is mediated by stabilization of the RyR1-FKBP12 complex, thereby reducing SR Ca^2+^ leak and restoring proper Ca^2+^ signaling. Notably, MP-001 also slowed the accumulation of reactive oxygen species (ROS), underscoring its dual role as a RyR1 stabilizer and antioxidant compound. Additionally, our data suggest that reshaping RyR1 function has a greater impact on muscle performance than the antioxidant effect alone, highlighting the centrality of RyR1-mediated Ca^2+^ leak in DISM.

In summary, our data support a model in which DOX-induced skeletal muscle dysfunction arises from two primary mechanisms, as shown in Figure 6: (a) direct effect on RyR1, likely caused by the destabilization of the RyR1-FKBP12 complex, leading to a decreased CDI and SR Ca^2+^ leak, which will induce an increased SOCE activity, and thus a rise in basal [Ca^2+^]_cyto_. Subsequently, this elevated cytosolic Ca^2+^ will indirectly promote mitochondrial ROS production, and (b) a direct DOX-induced mitochondrial ROS production, which likely co-occurs with its effect on RyR1. This overproduction of ROS further oxidizes RyR1, perpetuating Ca^2+^ dysregulation. This vicious cycle ultimately compromises muscle performance; however, it can be interrupted by targeted pharmacological intervention with a RyR1-FKBP12 reshaper at the level of the Ca^2+^ channel.

**Figure 6.**
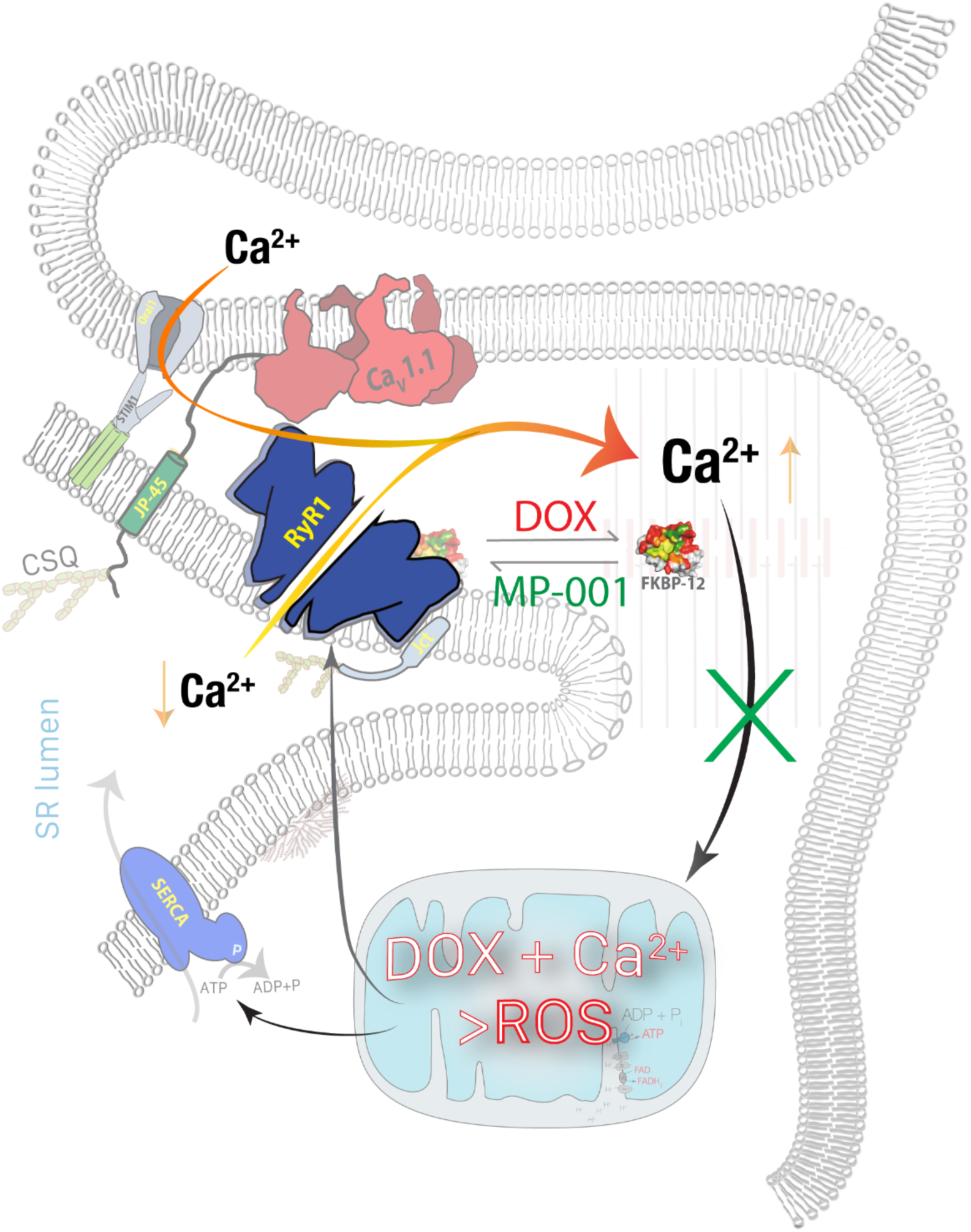
Simplified diagram with relevant proteins affected by DOX. DISM has multiple causes inside muscle fibers. DOX interferes with the regular function of mitochondria, leading to the production of ROS. Additionally, DISM is triggered by DOX-driven resting RyR1 function, which affects its regular activity and causes it to become leaky. The decrease in [Ca^2+^]_SR_ activates store-operated Ca entry (SOCE), increasing the basal [Ca^2+^]_cyto_ to abnormal levels. This last leads to increased oxidative stress, mitochondrial damage, and muscle dysfunction. This vicious cycle can be stopped, or at least reduced, by RyR1-directed inhibition using MP compounds, which stabilize RyR1 under DOX and oxidative stress conditions, becoming an effective anti-DISM therapeutic strategy.

These findings have important implications for the development of adjunct therapies to mitigate chemotherapy-induced muscle toxicity.

