## Supplemental Figures for "Disrupting the Feed-Forward Cycle of RyR1 Ca^2+^ Leak and Oxidative Stress Mitigates Doxorubicin-Induced Skeletal Myopathy"

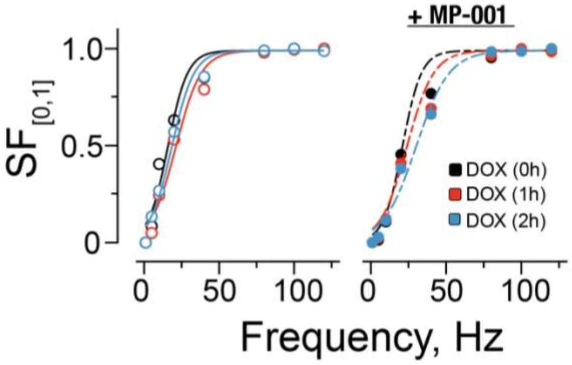


Figure S1. **Lack of kinetic variations in specific force signals in FDB muscles exposed to DOX.** Summary of maximum peak force values obtained at different frequencies of stimulation and at 1 and 2 hours after applying DOX and DOX + MP-001, normalized to the maximum value acquired at time 0h. Signals were normalized, and sigmoidal functions were plotted using the average of the fitting parameters in an N = 4 muscles per condition. Signals were not statistically different.

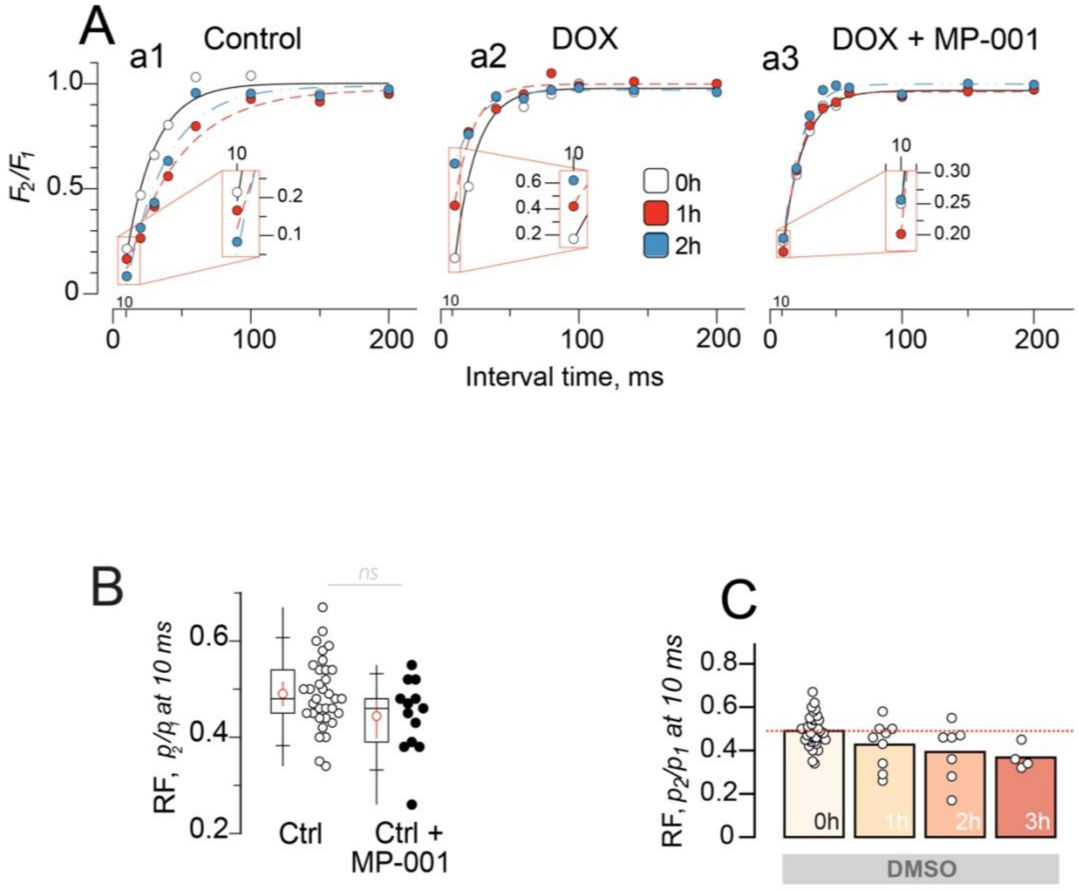


Figure S2. **SR Ca release elicited by two action potentials separated by interstimulus intervals of varying duration.** In ***A,*** an example of the fractional peak value (FP) of Ca-dependent fluorescence ($F_2/F_1$) is shown for each time interval and under the different conditions. In the inset of panels a1, a2 & a3, an example of FPs at 10 ms interpulse interval is shown. Values are significantly different at 10 ms when DOX is applied (*a2*), ***p*<0.01 in two-level hierarchical analysis. Panel ***B*** depicts the residual fraction (**RF**) calculated for controls with and without MP-001, showing no statistical differences. In ***C***, we show the natural decrease in the RF over time. ***p*<0.01 in two-level hierarchical analysis.

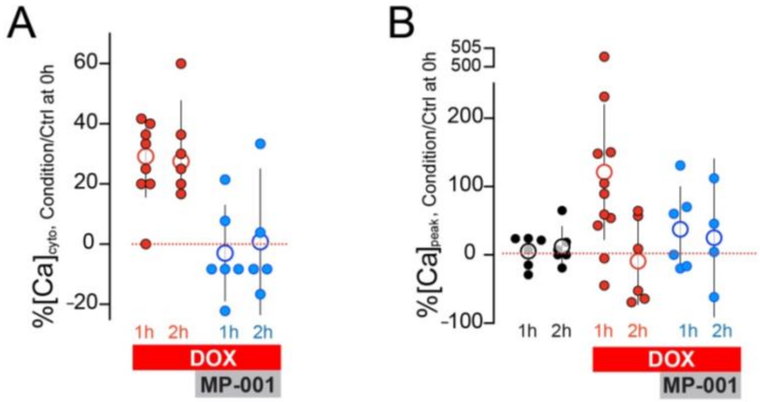


Figure S3. **Percentage of variation of basal and peak cytosolic Ca signals under DOX and MP-001 actions.** ***A*** summarizes the percentage of change in basal Ca with respect to basal conditions at 0h. Basal $[Ca]_{cyto}$ increases a 30% after 1 and 2 hours of DOX application (red symbols), and did not change in the presence of MP-001 (blue symbols). In panel ***B,*** we show the summary of peak [Ca] before and after DOX and DOX + MP-001 application. Values increase to more than 100% after the first hour of DOX application compared to the control at time 0h. Values are mean ± SD, *ns* represents not significant, ***$p<0.001$, in a two-level hierarchical analysis.
